# High-density spatial transcriptomics arrays for *in situ* tissue profiling

**DOI:** 10.1101/563338

**Authors:** Sanja Vickovic, Gökcen Eraslan, Fredrik Salmén, Johanna Klughammer, Linnea Stenbeck, Tarmo Äijö, Richard Bonneau, Ludvig Bergenstråhle, José Fernandéz Navarro, Joshua Gould, Mostafa Ronaghi, Jonas Frisén, Joakim Lundeberg, Aviv Regev, Patrik L Ståhl

## Abstract

Tissue function relies on the precise spatial organization of cells characterized by distinct molecular profiles. Single-cell RNA-Seq captures molecular profiles but not spatial organization. Conversely, spatial profiling assays to date have lacked global transcriptome information, throughput or single-cell resolution. Here, we develop High-Density Spatial Transcriptomics (HDST), a method for RNA-Seq at high spatial resolution. Spatially barcoded reverse transcription oligonucleotides are coupled to beads that are randomly deposited into tightly packed individual microsized wells on a slide. The position of each bead is decoded with sequential hybridization using complementary oligonucleotides providing a unique bead-specific spatial address. We then capture, and spatially *in situ* barcode, RNA from the histological tissue sections placed on the HDST array. HDST recovers hundreds of thousands of transcript-coupled spatial barcodes per experiment at 2 μm resolution. We demonstrate HDST in the mouse brain, use it to resolve spatial expression patterns and cell types, and show how to combine it with histological stains to relate expression patterns to tissue architecture and anatomy. HDST opens the way to spatial analysis of tissues at high resolution.

---

Tissues are composed of diverse cells organized to interact and execute complex functions. Charting the spatial organization of cells along with their molecular features provides a powerful lens into tissue function and is a cornerstone of disease pathology^1^. For example, understanding the spatial connections between different cells in the central nervous system (CNS) is critical for both basic brain function and the pathophysiology of neurodevelopmental disorders, neurodegenerative disease and brain tumors^2–4^. In recent years, methods for both single-cell and spatial profiling have advanced dramatically^1^, especially those focused on the transcriptome. In particular, massively parallel single-cell RNA-Seq (scRNA-Seq)^5,6^ can profile hundreds of thousands of dissociated individual cells, but does not directly recover the spatial position of those cells. In addition, it requires dissociation and cell manipulation, which can be difficult to perform; may require fresh tissue; and can introduce biases that lead to an altered molecular state or incomplete recovery of the tissue’s cellular constituents^7^. Conversely, spatial profiling approaches capture detailed spatial information in intact frozen or fixed tissue. However, current methods either require pre-selected limited marker signatures, are challenging to scale and apply to different biological systems, rely on non-standard instrumentation^8–13^ or have limited spatial resolution. In particular, Spatial Transcriptomics^14^ (ST) is a spatially barcoded RNA-Seq method that provides transcriptome-wide coverage in a variety of systems and can be applied to frozen sections on a glass slide, but its resolution is limited to 100 μm (on average 3-30 cells).

To bridge this gap, we developed High-Density Spatial Transcriptomics (HDST) and demonstrate its application to spatially profile large tissue areas in the mouse brain *in situ* at high resolution (Fig. 1a). In HDST, we deposit barcoded poly(d)T oligonucleotides with a randomly ordered bead array-based fabrication process. Specifically, beads carrying spatial barcodes are packed efficiently into wells^15^ and decoded prior to the biological experiment by a sequential hybridization and error-correcting strategy^15,16^. After a tissue section is placed on the slide, stained and imaged, RNA is captured and then profiled by RNA-Seq. This allows us to capture and reliably quantify transcriptomes in 2 μm features, while maintaining the histological two-dimensional tissue information intact.

**Figure 1.**
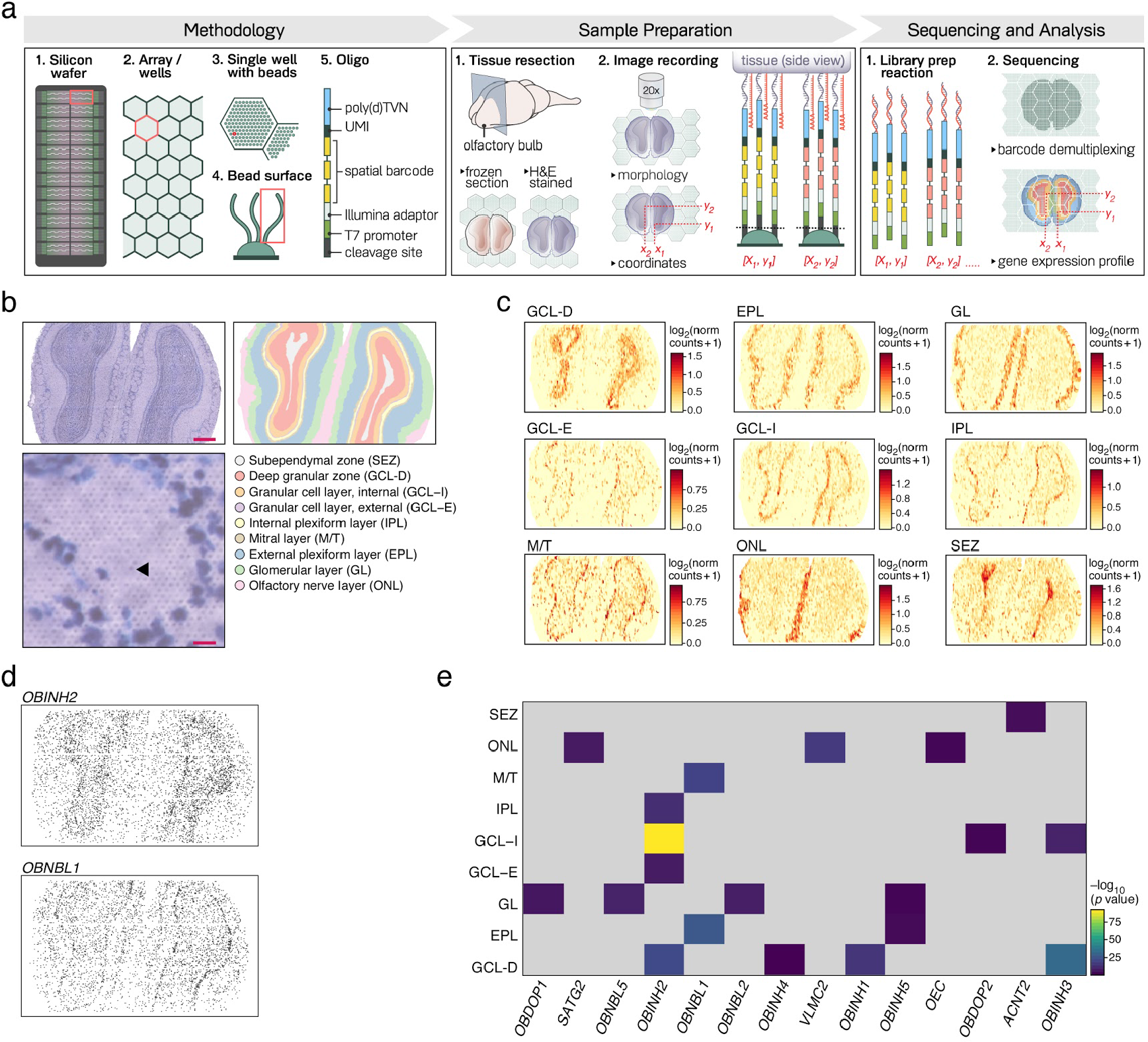
High-density spatial transcriptomics (HDST). **(a)** HDST workflow. Methodology: Barcoded beads are randomly deposited into single wells. The barcode carries a spatial oligonucleotide sequence encoding an (x,y) position in each individual well and array present on the silicon wafer. The complete oligonucleotide sequence then carries primers for mRNA capture. Sample preparation: Frozen tissue sections are placed on the HDST array surface, H&E stained and imaged for both morphology and recording the relative positions of each well with a barcoded bead to the tissue section. mRNA is captured on the oligonucleotide capture sequence and cDNA made. Spatial oligonucleotide sequence is covalently linked to the mRNA information for each cell in the tissue. Sequencing and Analysis: Standard pair-end sequencing libraries are made, spatial barcodes demultiplexed and the whole tissue section profiled on a transcriptome-wide scale with HDST. **(b)** Labeling of morphological layers. HDST H&E image of a main olfactory bulb (upper left; scale bar 300 μm) and matching HDST (x,y) barcodes annotated into 9 different morphological areas (upper right, color code). Zoomed-in view of a glomerulus structure (lower left; scale bar 10 μm) with a single HDST bead-well denoted with an arrowhead. **(c)** Layer-specific differential gene expression patterns in HDST. Shown is the summed normalized expression of positively enriched signature genes (**Supplementary Table 2**) significantly (FDR<0.1, two-sided *t*-test) associated with each layer as annotated in (b). **(d)** Lightly binned spatial barcodes assigned (black) to each of two cell types (OBINH2: inhibitory neurons, OBNBL1: neuroblasts). **(e)** Enrichment of lightly binned spatial barcodes with assigned cell types (columns) to morphological layers (rows). −*log*_10_(p-value) (Fisher’s exact test, Bonferroni adjusted, p-value < 0.01) represents the colorbar. Grey tiles represent non-significant values.

To produce a high-resolution and high-density bead array, we generated barcoded beads with a split-and-pool approach, randomly placed them into an array of >1.4M 2 μm wells, and then decoded which bead is in which well (Fig. 1a, **Methods**). The resulting beads had sufficient barcode complexity to avoid substantial redundancies in duplicate spatial (x,y) locations. For example, of 2,839,865 individual beads we expect that that 69.5% of the beads would be uniquely useable after decoding. We fabricated the bead-in-well slide array in a 1,918×765 matrix for a total of 1,467,270 wells arranged into a hexagonal pattern. We estimated the well size at 2 μm spaced at 3 μm center-to-center. The bead-in-well approach prevents barcode mixing between individual wells and the hexagonal design enables maximum tissue capture. To give a spatial address to each barcode and bead, we decode the barcode sequences with several rounds of sequential hybridization^16^. In each round, we hybridize a set of complementary and labeled decoder oligonucleotides (“decoders”), record fluorescence across the entire slide area, and then dehybridize the decoders. We repeat this hybridization and recording process with the same decoder sequences now labeled with a different fluorophore and finally with a set of unlabeled decoders. The process was repeated for *log*_3_*N*times (13 times for the array presented here), with *N* representing the number of sequences to be decoded and 3 being the number of labels (green, red and “dark”) used. Each bead and spatial barcode is now encoded with a unique spatial color address^16^ creating a HDST array. Arrays are decoded and images processed in a production setting with ~3h total processing time. Following decoding, we capture, barcode and profile RNA (Fig. 1a). Tissue sections placed onto the bead array surface are stained and imaged. We then gently permeabilize the tissue to capture mRNA molecules onto the respective bead capture oligonucleotides, leading to effective direct *in situ* barcoding. Next, we perform a reverse transcription reaction, library preparation and RNA-Seq (**Methods**). The resulting HDST experiment provides a 1,400-fold increase in resolution compared to previous ST technology while still spatially profiling same-sized tissue sections.

To test HDST, we profiled the main olfactory bulb (MOB) of the mouse brain. Neurons in the MOB have traditionally been defined by the presence of neuronal cell bodies in the different morphological layers^17^. We thus sought to see whether molecular data from HDST can be related to layer and other histological data. We analyzed three replicate sections by HDST, and included a standard ST^14^ dataset from the same brain region for comparison. We then tested the performance of HDST in two key tasks: (**1**) the generation of high-resolution spatial expression patterns of individual genes, and (**2**) the detection of cell types and their assignment to correct, high-resolution positions.

First, we confirmed that RNA capture was specific and in overall agreement with bulk RNA-Seq controls, despite the relatively low number of transcripts captured per spatial barcode. After accounting for barcode redundancy (“clashing”), decoding efficiency and stringent barcode demultiplexing (Supplementary Fig. 1a**, Supplementary Table 1**), we proceeded to analyze the spatial RNA-Seq profiles. At saturating sequencing depth (Supplementary Fig. 1b), 81.5±1.8% of all genes detected were located within the area physically covered by the tissue specimen (without using any lower cutoffs), with almost 140,000 barcodes (Supplementary Fig. 1c) generating spatial profiles per assay (n=3). Although the average number of UMIs per barcode location was low (Supplementary Fig. 1d), a we detected a very distinct spatial *in situ* tissue profile following the tissue boundary (Supplementary Fig. 1-e), suggesting the specificity of our detection. Moreover, a combined “bulk” expression profile of each HDST dataset correlated significantly with published MOB bulk RNA-Seq (Supplementary Fig. 2a; Spearman’s rS = 0.69±0.02; mean±sd) and across the three replicate experiments (Spearman’s rs = 0.82±0.06; mean±sd). Most of the detected genes agreed between the bulk and HDST datasets (Supplementary Fig. 2b).

For the first task of spatial expression profiles, we successfully used HDST data in supervised analysis to correctly identify layer-specific expression signatures. To this end, we first annotated morphological layers (**Methods**) from the H&E stain of each specimen (Fig. 1b). We further reasoned that two closest proximal beads in each direction, for a total of 24 neighbor positions (r ~ 6.5 μm), are likely to capture transcripts from the same cell and layer. We thus enhanced the signal by light binning, through pooling of reads within a short range (*e.g.*, 5X compared to the 5×5 hexagonal wells). On average, each bin had observations from 3.5±1.9 (mean±sd) (x,y) beads and 10.7±9.1 (mean±sd) overall UMIs. Finally, we assigned each “enhanced” bin to a layer, to robustly identify differentially expressed (DE) genes between the morphological layers (**Methods**). Following a smoothing Gaussian filter on the binned data (16.9±11.3 (mean±sd) UMIs per bin), we performed a two-sided *t*-test (FDR<0.1), identifying DE signatures specific to morphological layers (Fig. 1c, Supplementary Table 2, Supplementary Fig. 3a-c). We confirmed that layer-enriched upregulated DE genes (FDR<0.1; *log*_2_ fold change>1.5) were correctly assigned by comparison to their *in situ* hybridization (ISH) score in the Allen Brain Atlas (ABA)^18^ coronal dataset, both as average signatures (Supplementary Fig. 4a) and by comparison of individual patterns (Supplementary Fig. 4b).

To test cell type spatial assignment, we developed a multinomial naive Bayes classifier to map the high-resolution, but sparse, spatial data in HDST to cell type annotations by integration with scRNA-Seq (**Methods**). We first used scRNA-Seq UMI counts^19^ to compute the maximum likelihood estimates of the multinomial parameters for each cell type (i.e. normalized mean gene expression) (**Supplementary Table 3**). We then estimated the likelihood that a given HDST gene expression profile originated from each of the scRNA-Seq cell types and using posterior probabilities assigned cell types to the barcode locations (**Methods**, **Supplementary Table 4**).

Approximately 30% of spatially barcoded HDST profiles were confidently assigned to a single cell type. To estimate the sensitivity of our cell assignment to read depth and spatial resolution, we again decreased the resolution of HDST data using binning (**Methods**), with the 5X mimicking a single cell (“sc-like”) and 38X resolution mimicking ST data (“ST-like”). We repeated the cell type assignment task using the 5X, 38X and ST data^14^ (Supplementary Fig. 5a). The posterior probabilities of the cell type assignments increased in the larger bins (Supplementary Fig. 5b-c**, Supplementary Table 5**) and a single cell type could be confidently predicted in 69.14% of all barcoded (x,y) positions in the lightly binned “sc-like” (5X) data (Supplementary Fig. 5d), with differential markers driving the assignment task (**Methods**, Supplementary Fig. 5e**, Supplementary Table 6**). In comparison, only 0.37% of (x,y) positions could be assigned to a single cell type in ST using this approach.

Finally, we coupled the spatially assigned cell types to the morphological layers (Fig. 1d, **Methods**). Some populations exhibited layer-specific patterns (p-value<0.01, Fisher’s exact test with Bonferroni adjustment; Fig. 1e). In total, we recovered cells from 14 cell types (out of 63 tested ones) as layer enriched, spanning 10 neuronal, 3 glial and one vascular population. For example, a neuroblast population (OBNLB1) was identified as enriched in the mitral (M/T) and external (EPL) plexiform layers, the deep granular zone (GCL-D) was enriched with inhibitory neurons (OBINH1-4), the subependymal zone (SEZ) with non-telencephalon astrocytes (ACNT1) and the olfactory nerve layer (ONL) with olfactory ensheathing cells (OEC), vascular cells (VLMC2) and satellite glia (SATG2). Different neuronal populations (GABAergic; OBNBL5 and OBINH5, dopaminergic periglomerular; OBDOP1 and VGLUT1/2 OBNLB2 neuroblasts) were found in the glomerular layer (GL). Many of these populations have previously been reported to be associated within the same specific morphological layers^19^ with inhibitory neurons (OBINH2) dominting granular (GCL-E, GCL-I and GCL-D) and the adjacent internal plexiform (IPL) layers. Our results corroborate traditional classification of neurons^17^ in the main olfactory bulb. Further development of computational methods for modeling cell types using HDST data and their explicit integration with image analyses will likely improve our ability to both detect and observe deviations from canonical (or previously observed) cell types.

We show that HDST is a robust high-resolution approach providing *in situ* spatial information on tissue composition. The technology relies on robust and commoditized tissue, molecular, bead-array and imaging modular tasks, which can be readily deployed across the scientific community. HDST uses standard histological stains, providing the means to relate morphology, extracellular features and gene expression. While the data is relatively sparse in this first demonstration of the technology, it is highly specific, and can be interpreted by computational integration with morphological features and single-cell profiles. Further development of HDST and associated computational methods will help us understand tissue organization and function in health and disease.

## Methods

### Bead production

We used a split-and-pool approach to generate a total of 2,893,865 different quality controlled 2μm silica beads. A primer precursor containing (1) the T7 promoter, (2) an Illumina sequencing handle followed by (3) the first 15 bp “Spatial pool” and (4) a 7 bp “bridge” oligonucleotide sequence /AmC6/UUUUUGACTCGTAATACGACTCACTATAGGGACACGACGCTCTTCCGATCT-Spatial_barcode_Pool1-Bridge1) (IDT) were linked to the bead surface using amine chemistry^16^. After linkage, in order to increase the bead pool size (defined as the number of uniquely barcoded beads in the pool), we pursued two additional sequential ligation steps, adding 14 bp and 15 bp pools of spatial barcode sequences (with similar GC content and Tm), using two different bridge oligonucleotide sequence pairs (with /Pho/Bridge2-Spatial_barcode_Pool2-Bridge3 being ligated to Bridge1 through a complementary Bridge1‘Bridge2’ 14 bp sequence). We relied on double-stranded ligation using T4 DNA ligase (NEB) with the ligation oligonucleotide added in a 2:1 ratio to the precursor oligo sequence and following the manufacturer’s ligation protocol. In the second ligation step, the newly ligated sequence ending with the Bridge3 sequence acted as the precursor for the next spatial pool (/Pho/Bridge4-Spatial_barcode_Pool3 ligated through Bridge3’Bridge4’). In this final ligation step, the last barcode sequence was followed by a 7 bp unique molecular identifier and a stretch of 20 (d)Ts and VN to ensure efficient mRNA capture on the surface. The ligated Bridge1Bridge2 sequences read GACTTGTCTAGAGC and Bridge3Bridge4 TGATGCCACACTACTC. All sequences except the first precursor oligonucleotide containing the Spatial_Pool1 were synthesized on Illumina’s “Big Bird” oligonucleotide synthesis platform.

### Array generation

The complete bead pool was used to load a total of 1,467,270 hexagonal well positions covering a 13.7 mm^2^ area (5.7 mm x 2.4 mm). The wells were etched in a planar silica slide and polished to 1 μm height. A total of 24 such areas were made on each slide. The bead pool was loaded in ethanol onto the planar slides with shaking so the beads would easily be captured into well and unused beads washed away.

### Array decoding

Two sets of complementary and fluorescently labeled (FAM and Cy3) oligonucleotides were synthesized, deprotected and purified. An additional set of unlabeled but still complementary probes were made to enable error-checking. Each set represented an individual decoder pool (10nM). Loading, hybridization, fluorescence detection and probe stripping were performed as described previously^16^ and using Illumina’s IScan system where each channel (FAM; green or Cy3;red) was imaged separately. Each (x,y) position was encoded with a unique combination of three colors (FAM; Cy3 and “dark”). Raw decoded spatial arrays and corresponding decoder files were shared by Illumina after bead array production in the standard Illumina DMap format. Barcode decoding (including empty wells) and redundancy percentages based on the Illumina decoding process were calculated and reported in **Supplementary Table 1**.

### Tissue samples

Adult C57BL/6J mice (at 12 weeks age) were euthanized and their main olfactory bulb dissected. The samples were then frozen in an isopentane (Sigma-Aldrich) bath kept at −40°C, and transferred to −80°C for storage until sectioning. The frozen bulbs were embedded at −20°C in Tissue-Tek OCT (Sakura) compound. Cryosections were taken at 10 μm thickness and deposited on prechilled slides containing barcoded arrays.

### Tissue staining and imaging

Tissue sections were first adhered to the surface by keeping the slide at 37°C for 1 min. Immediately after, a fixation step on the slide surface was performed using 4% neutral buffered formaldehyde (Sigma-Aldrich) in 1x phosphate buffered saline (PBS, pH 7.4) for 10 min at room temperature (RT). The slides were then washed once in PBS to ensure proper formaldehyde removal. The sections were stained using standard hematoxylin and eosin (H&E) staining, as previously described^14^, and imaged with a Ti-7 Nikon Eclipse with a NB filter in fluorescent mode to expose the samples to a bright field light source and the reflections collected on a color camera. This allowed histological imaging of a dark slide on a standard epifluorescence microscope.

### RNA-Seq library preparation and sequencing

We followed a protocol as described in detail in Stahl *et al*^*14*^ and Salmén *et al*^*20*^. Briefly, tissue sections were gently permeabilized using exonuclease I buffer (NEB) and pepsin, followed by *in situ* cDNA synthesis overnight at 42°C using Superscript III (ThermoFisher Scientific) supplemented with RnaseOUT (ThermoFisher Scientific), such that the transcript was reverse transcribed into cDNA and covalently linked to the spatial barcode. Tissue sections were then digested using proteinase K (Qiagen) and the barcode information cleaved using a Uracil-Specific Excision Reagent (NEB) targeting the 5d(U) stretch at the 5’ end on the barcoded oligonucleotides. The collected material was then processed into libraries as described in Jemt *et al*^*21*^. The libraries were sequenced on a Illumina Nextseq 500 instrument with v2 chemistry with paired-end 300 bp reads (R1 125 bp and R2 175 bp).

### HDST data pre-processing

FASTQ files were processed using the ST Pipeline v1.5.0 software^22^. The forward read contained both the barcode sequence and the bridge sequence used for the sequential ligation steps. Bridge sequences were trimmed and removed prior to any barcode mapping. Forward reads were trimmed retaining the following bases: 1-15, 31-45 and 61-76. This created a trimmed forward read containing a spatial barcode followed by a UMI. Transcripts (only in reverse read) were mapped with STAR^23^ to the mm10 mouse reference genome. Mapped reads were counted using the HTseq count tool^24^. Spatial barcodes were demultiplexed using an optimized implementation of TagGD^25^ as described^22^. We allowed a 2 bp “padding” overhang on the total length of the spatial barcode. This enabled correction for either insertion or deletion error at the very beginning (1 bp) or the very end (1 bp) of the barcode sequence. Demultiplexing was based on building a hashmap of 11 bp *k*-mers. All barcodes were then compared to the complete barcode sequence allowing a Hamming distance of 4 mismatches. Start position for the UMI was 77 and end position 82 in the forward read. UMI duplicated sequences paired to the annotated reads were collapsed using a hierarchical clustering approach. All UMIs were clustered using the spatial barcode, mapped gene locus (with an window offset of 250 bp to account for alternate termination sites^24,26^) and strand information. Only one mismatch error was allowed in UMI clustering. This process then generated a bead-by-gene count matrix with a Cartesian (x,y) coordinate assigned with gene expression information.

### Histological image processing

To relate the histological image and the counts matrix, we assigned image pixel coordinates to the centroids of each bead well^27^. This ensured proper alignment of tissue boundaries in the image and allowed us to select barcodes that were located spatially underneath tissue boundaries. We took the same approach to detect the arrays’ boundaries and corners, and assumed a perfect hexagonal well matrix, which is reasonable given standardized production and quality control specifications for each slide. We translated pixel coordinates into fixed centroid (x,y) coordinates using the total detected area of the array. The coordinate names were then matched to the barcode decoder file used in the HDST pre-processing step.

### Image annotation

Images used in the study were annotated with an interactive user interface for selecting spatial barcodes and their (x,y) coordinates based on the tissue morphology. Each (x,y) barcode position could be assigned to one or more of the 9 distinct regions in the mouse olfactory bulb: Olfactory Nerve Layer (ONL), Granular Cell Layer External (GCL-E), Granular Cell Layer Internal (GCL-I), Deep Granular Zone (GCL-D), External Plexiform Layer (EPL), Mitral Layer (M/T), Internal Plexiform Layer (IPL), Subependymal Zone (SEZ) and the Granular Cell Layer (GL). For HDST, exactly one of the nine regional tags was assigned to one (x,y) spatial barcode. For ST, more than one tag could be assigned per (x,y) spatial spot location, when one (x,y) standard ST spot area spanned more than one layer.

### Binning of HDST data

We divided the total area of each HDST array into non-overlapping bins, each covering an area of *X×X* beads, where *X∈{5,38}*, and summed the UMIs of beads within each spatial bin. In order to ensure appropriate bin sizes, we first considered all manufactured wells as a 1918×765 matrix. On average, around 1,370 (x,y) wells filled with beads would size up to one ST spot (100 μm; *X* = 38) when taking into account the center-to-center distance between two wells. We used this as our maximum *X*. We took the binned data containing 1,370 wells per bin and took every second bin into account in both x and y directions. This was to ensure space between two ST spots would be accounted for. This bin size was called “ST-like” in all further analyses. We did not take into consideration that this bin actually represents 63% of the transcriptome profiled per ST spot due to the space between two hexagonal wells. Second, we made bins with fewer wells per bin in a logarithmic manner until reaching the smallest bin (5X). The 5X bin was referred to as single cell like of “sc-like”.

### Spatial differential expression analysis

Binned 5X data was smoothed using a Gaussian kernel (5×5) with 0.5 standard deviations equally in both x and y directions. The smoothed binned data was then scaled such that the maximum expression value stayed the same. Smoothed HDST data was normalized to an average UMI count per bin. We performed a two-sided t-test (FDR<0.1) to identify DE genes for each HDST morphological region. The top 500 genes identified per morphological layer with a *log*_2_ fold change > 1.5 (one vs. rest) were identified as differentially expressed and used in further analyses (**Supplementary Table 2**).

### Validation of differentially expressed genes

To validate layer-specific gene expression in the HDST data, we performed enrichment analysis using layer specific gene sets from the Allen Brain Atlas as reference. Layers annotated in both datasets were used in the analysis with all HDST granular layers merged into one instance to be comparable to the data provided in the Allen Brain Atlas. Genes with a layer specific *log*_2_ fold change of greater than 1.5 (implying upregulation) and FDR<10% were tested for enrichment in the layer-specific gene sets (“expression fold” change greater than 1.5) from the Allen Brain Atlas. Only genes passing the respective fold-change thresholds in both data-sets were included in the analysis. Images for the top gene present in each layer were downloaded from ABA’s *High Resolution Image Viewer* and stitched using Fiji^28^.

### Cell type assignment to HDST barcodes

We downloaded the matrix containing mean expression values *x* per cell type *j* from Zeisel *et al*^*19*^ using the loompy package (https://github.com/linnarsson-lab/loompy). We subset the matrix to contain only cell types annotated in the olfactory bulb (OB) and non-neuronal cell types for a total of 63 cell types (**Supplementary Table 3**). The vector of probabilities *Θ*_*j*_ of each gene being captured in a cell type *j* is defined as follows:

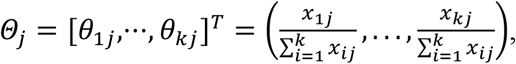

where *i* = *gene*, *j* = *cell type*, *k* = *total number of genes*, *x* = *mean expression*

We calculate the likelihood *L* of the cell type *j* specified by *Θ*_*j*_, given the observed UMIs *b* per gene for a HDST (x,y) barcode as follows:

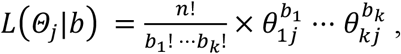

where *b* = *vector of UMI counts per gene for an individual HDST (x, y) profile*

*n* = *total number of UMIs for an individual HDST (x,y) profile*

For each HDST (x,y) and cell type *j*, we calculated the ratio between the likelihood of that cell type *L*_*j*_ and the likelihood of the most likely cell type *L*_*max*_ as a measure to assess how good the secondary cell type assignments are compared to the most likely cell type (primary assignment):

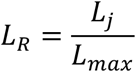

For each HDST (x,y) and cell type *j*, we calculated the posterior probability i.e. normalized likelihood *L*_*N*_ as the ratio between the likelihood of that cell type *L*_*j*_ and the sum of all likelihoods for that barcode *L*_*tot*_ as a measure to assess how distinct each assignment is compared to all others:

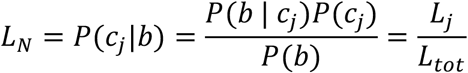

where *P*(*c*_*j*_) denotes the uniform prior for cell type *j*. *P*(*c*_*j*_|*b*) and *P*(*b*|*c*_*j*_) represent the posterior probability and the likelihood, respectively. *P*(*b*), the evidence term, is defined as 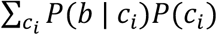 and used for the normalization.

Finally, to test against the null-hypothesis that HDST (x,y) expression profiles originate from random expression profiles for each cell type *j* and respective *Θ*_*j*_, we shuffled each *Θ*_*j*_ in 10,000 iterations while keeping the distribution of UMIs *b* as in the corresponding HDST (x,y) expression profile. We then calculated the randomized likelihood *L*_*R*_ for each HDST (x,y), cell type and iteration. Finally, an empirical p-value *p*_*emp*_ was calculated for each HDST (x,y) and cell type assignment as the fraction of *L*_*R*_ that yielded a likelihood higher or equal to the cell type likelihood *L*_*j*_, with correction for multiple testing (Benjamini-Hochberg). For each HDST (x,y) the cell type with the highest likelihood (*L*_*R*_ = 1) was considered the primary assignment and all cell types with *L*_*R*_ ≥ 0.8 were considered secondary assignments. Finally, for a cell type assignment to be considered valid, we required *L*_*N*_ ≥ 0.1 and *P*_*emp*_ ≤ 0.01. For further analysis only HDST (x,y) with exactly one valid cell type assignment were considered. Cell type assignments, ratios and the empirical p-values for all HDST (x,y) profiles have been reported in **Supplementary Table 4**. HDST (x,y) positions that yielded multiple valid cell type assignments (*L*_*N*_ ≥ 0.1 and *P*_*emp*_ ≤ 0.01 and *L*_*R*_ ≥ 0.8) were also reported in **Supplementary Table 4**. Cell types assignments were repeated in the same fashion for all bin sizes and reported in **Supplementary Table 5**. Differential expression analysis was performed between the cell types using two-sided *t*-test (*log*_2_ fold change of greater than 1.5 and FDR<10%) and reported in **Supplementary Table 6**.

### Auxiliary data pre-processing

Public bulk RNA-Seq datasets^14^ were downloaded from NCBI’s SRA with accession PRJNA316587 and mapped to the mm10 reference and UMI filtered using the ST pipeline v1.3.1. Averaged and naively adjusted^29^ bulk gene expression signatures were compared to those of the three replicates created with HDST and normalized the same way. Allen Brain Atlas (ABA) gene lists were downloaded from the ABA API using the ConnectedServices module of the allensdk Python package version 0.16.0. The standard ST data as a counts matrix was downloaded from http://www.spatialtranscriptomicsresearch.org/.

## Code availability

All computer code has been deposited on GitHub.

## Data and materials availability

The data has been deposited to NCBI’s GEO archive and in the Single Cell Portal.

## Acknowledgments

We thank the NGI Stockholm and SciLifeLab for providing infrastructure support. We thank Leslie Gaffney and Ania Hupalowska for help with the preparation of figures. We thank Jennifer Rood for help with manuscript preparation. We acknowledge support from the Flatiron Institute and the Simons Foundation. Work was supported by the Knut and Alice Wallenberg Foundation and the Swedish Research Council (to S.V., J.L. and P.L.S.), the Klarman Cell Observatory and HHMI (to A.R.).

## Contributions

S.V., J.F., J.L. and P.L.S. designed the experiments. S.V., F.S. and L.S. performed the experiments. S.V., J.K. and A.R. designed the analysis approaches. G.E. and J.K. devised and conducted the analyses with help from T.Ä., R.B., L.B. and J.F.N. S.V. and J.G. implemented the graphical interface for the annotation tool. S.V., F.S., J.L and P.L.S. co-developed the high-density arrays with M.R. S.V. and A.R. wrote the manuscript with input from all the authors. All authors discussed the results.

## Tables

**Supplementary Table 1.** Decoding and redundancy report for all slides and arrays used in the custom pool bead array batch.

**Supplementary Table 2.** Differentially expressed genes between different morphological layers after light binning and smoothing.

**Supplementary Table 3.** List of mean gene expression values followed by the vector of probabilities for all cell types used in the analysis.

**Supplementary Table 4.** Cell type assignments per HDST (x,y) position.

**Supplementary Table 5.** Cell type assignments for different sized bins.

**Supplementary Table 6.** Differentially expressed genes between different cell types after light binning.

**Supplementary Figure 1.**
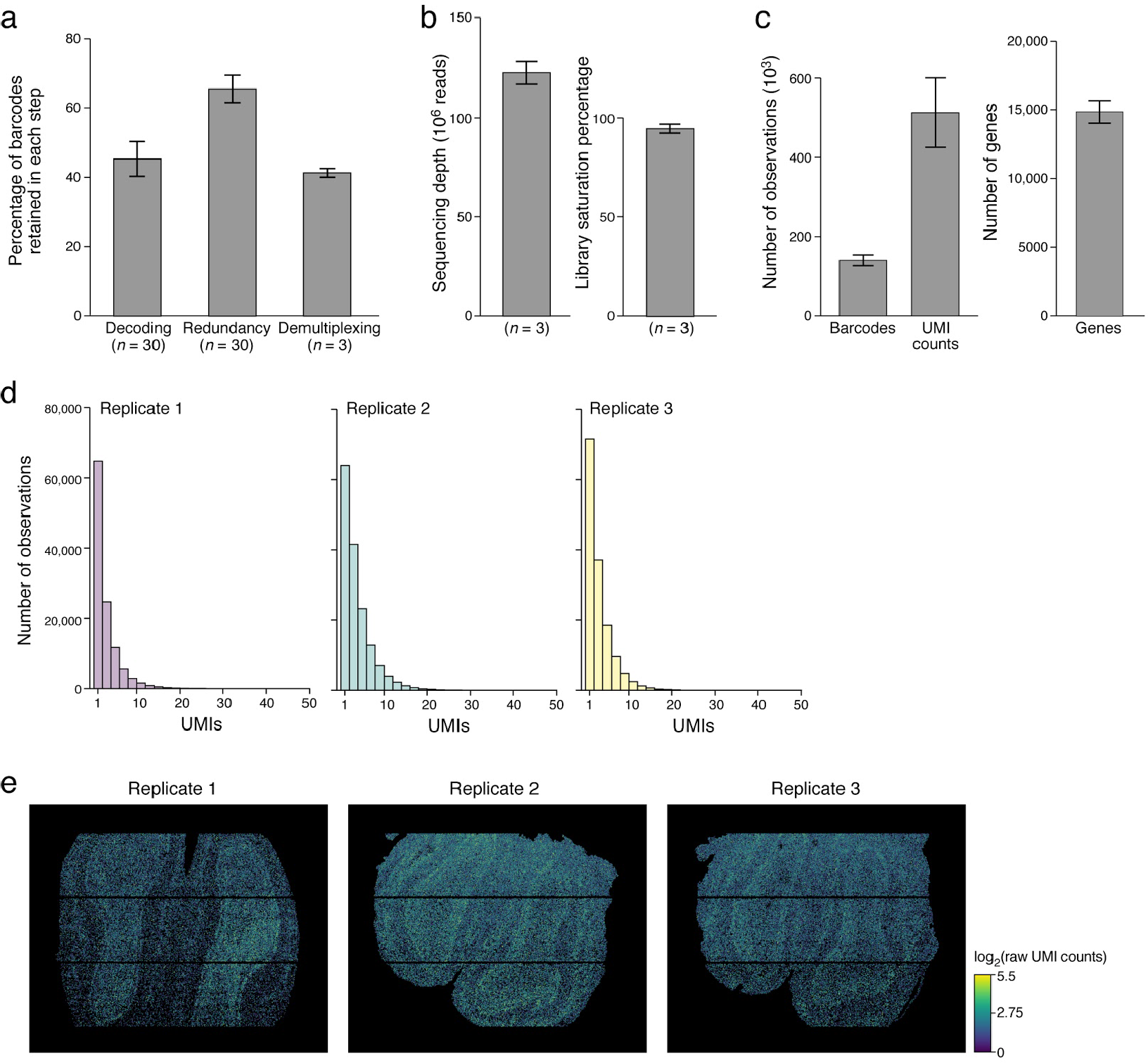
HDST array performance. **(a)** Barcode usage. Mean percent of spatial barcodes (y axis) that are retained in each of the following steps: after decoding (n=30 slides), removal of “clashing” (redundant) barcodes (n=30 slides) and barcode demultiplexing after sequencing (n=3). **(b)** Sequencing depth (y axis, left) and library saturation (y axis, right) for three replicate MOB HDST libraries (n=3). **(c)** Complexity of barcodes, UMIs and genes for three replicate MOB HDST libraries. Average number (y axis) of barcodes and UMI counts (left, x axis) and genes (right, x axis) recovered under the tissue boundaries after sequencing and filtering. **(d)** Recovered number of UMIs (x axis) per spatial barcode for each replicate library. **(e)** HDST specificity to the tissue section. Total UMIs per spatial barcode for all three MOB HDST replicates. Number of UMIs (log2(raw UMI counts), color bar) mapped at each position on the HDST array. Error bars: s.e.m. in all respective panels.

**Supplementary Figure 2.**
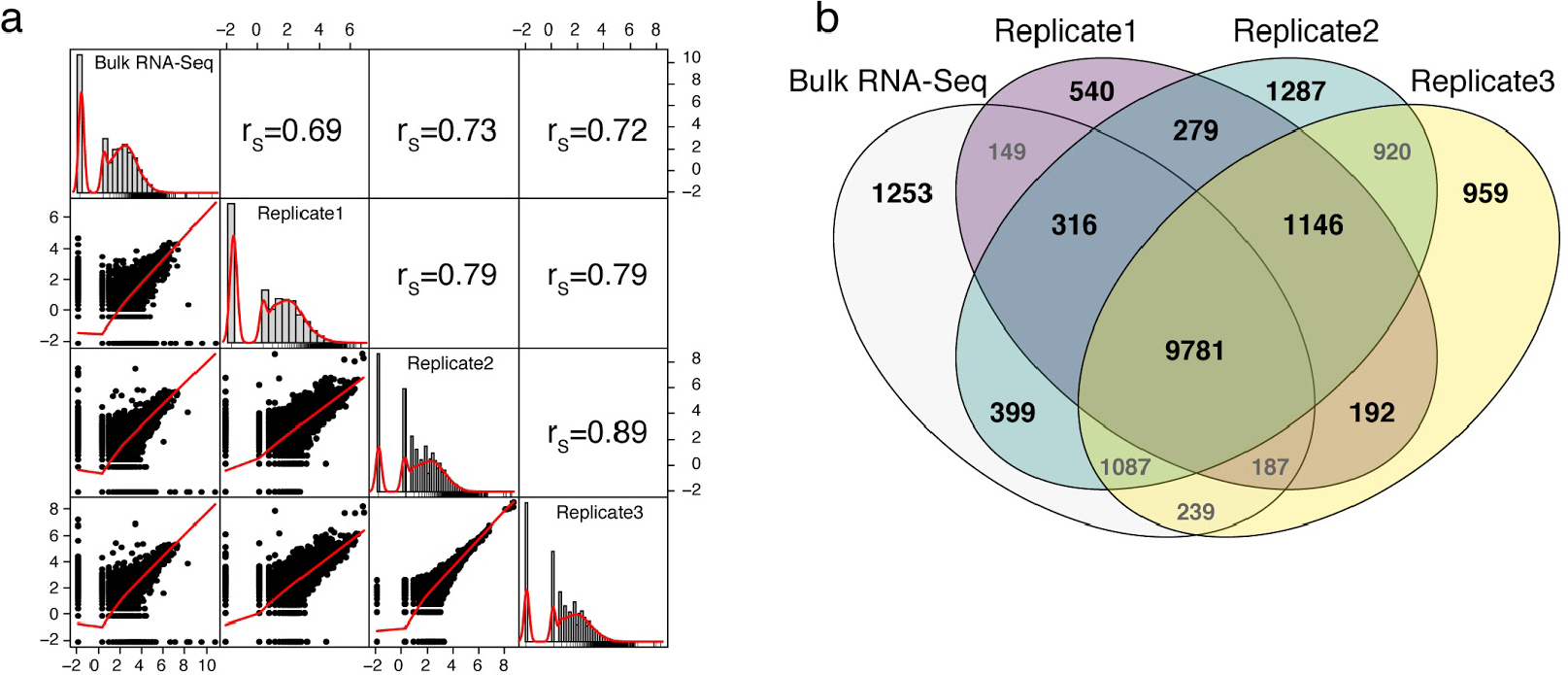
HDST agrees with bulk RNA-Seq. **(a)** Correlation of average gene expression profiles for each HDST replicate and for bulk RNA-Seq. Histogram of UMIs for each sample. Denoted are the Spearman correlation coefficients between groups. Red line represents fitted density distributions. **(b)** Numbers of genes detected in each of three HDST replicates and bulk RNA-Seq and their intersections.

**Supplementary Figure 3.**
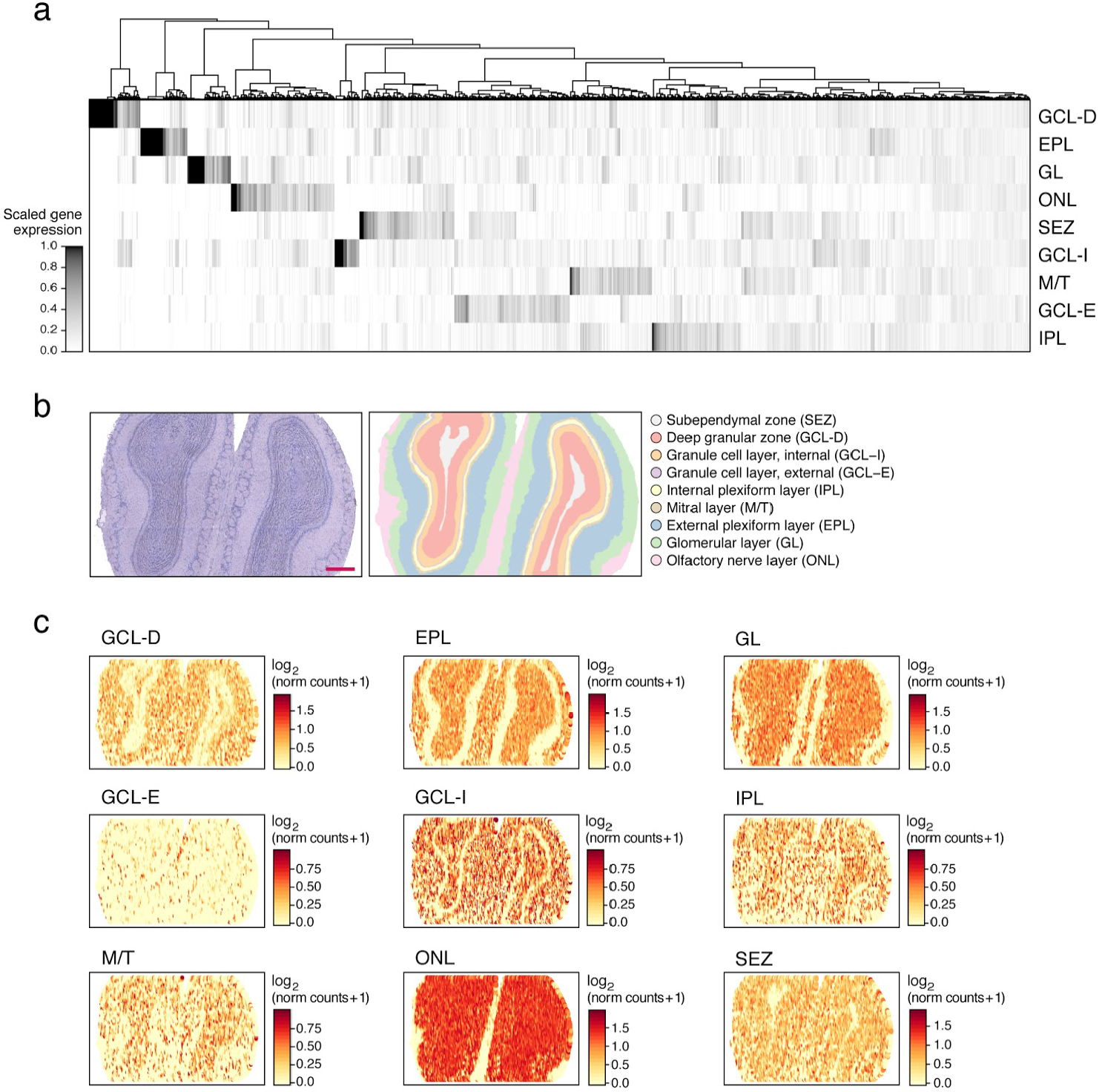
Layer-specific expression can be correctly recovered by HDST. **(a)** Differentially expressed genes between morphological layers in HDST. Shown is the scaled expression level (color bar, genes expression is scaled between 0 and 1 by subtracting the minimum across layers and dividing by the maximum) of genes (columns) that are recovered as differentially expressed between layers (rows) based on lightly binned and smoothed data. **(b)** Labeling of morphological layers. HDST H&E image of a main olfactory bulb (left; scale bar 300 μm) and matching HDST (x,y) barcodes annotated into 9 different morphological areas (right, color code). **(c)** Genes down regulated in specific layers. Shown is the spatial organization of barcodes, colored by summed gene expression for all genes differentially expressed in a specific morphological layer (label).

**Supplementary Figure 4.**
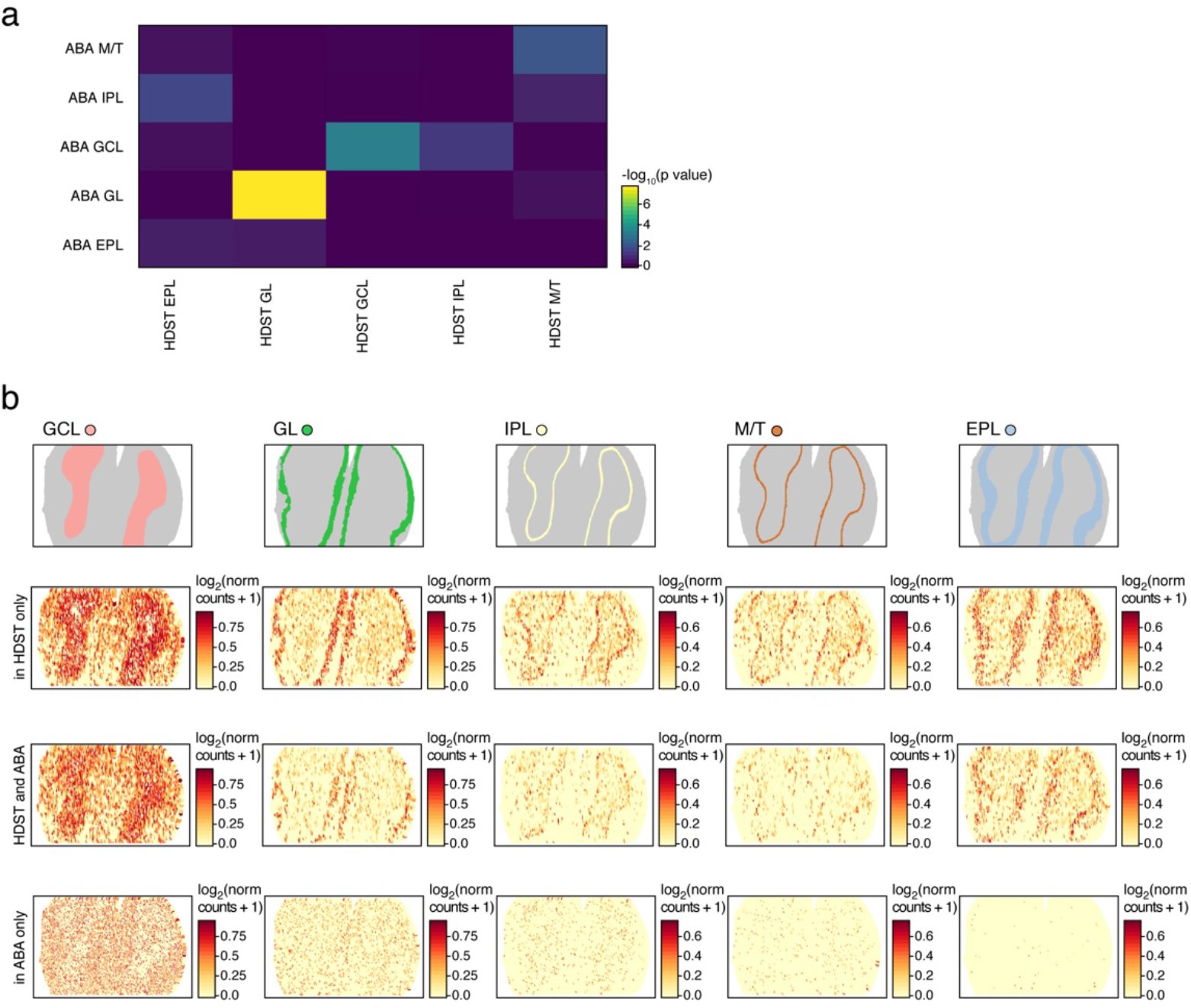
Spatial expression patterns from HDST are validated in the Allen Brain Atlas. **(a)** Layer specific signatures agree between HDST and the Allen Brain Atlas (ABA) for the major layers: granular (GL), glomerular (GCL) and mitral layers (M/T). −*log*_10_(p-value) of Fisher’s test (color scale) for enrichment of genes associated with each layer in lightly binned HDST (columns) and in the Allen Brain Atlas (rows). **(b)** Consistency in gene patterns. Shown are illustrative examples of summed gene expression (columns) whose spatial expression is found only as HDST specific, matches between HDST and the Allen Brain Atlas data or is found only in the Allen Brain Atlas (rows). Top illustration denotes layer-specific annotations (color code) in HDST where the grey color annotates all barcode positions not present in that layer.

**Supplementary Figure 5.**
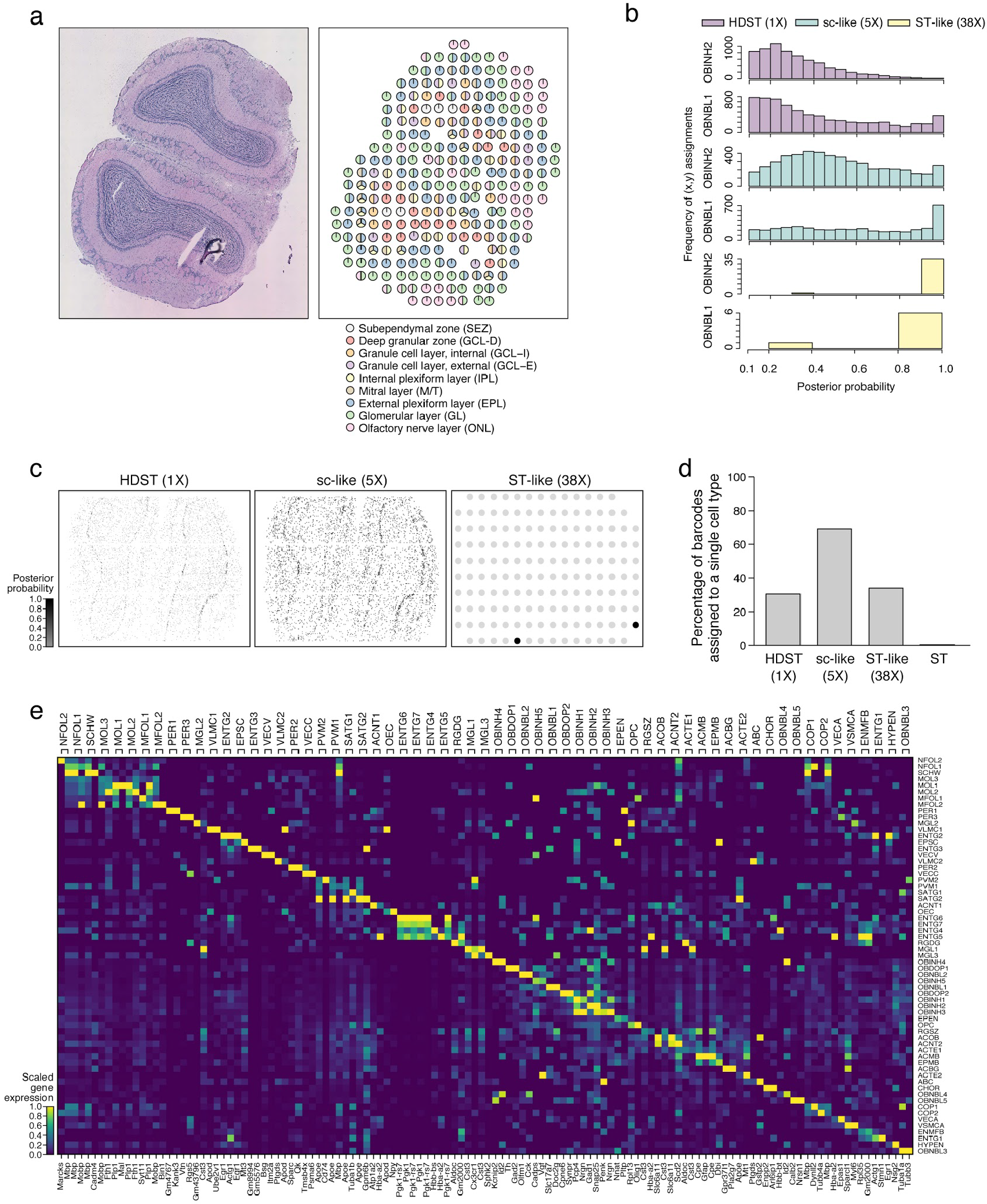
Light binning enhances HDST data cell typing. **(a)** H&E image of a main olfactory bulb (left), matching ST barcodes annotated into 9 different morphological areas (right, color code). **(b)** Impact of light “sc-like” (5X) and large (38X) “ST-like” binning on posterior probabilities of spatial barcode assignments to two of the selected cell types (OBINH2 and OBNBL1). Frequency distribution of cell type posterior probabilities when using binned (5X, cyan and 38X, yellow) and HDST (1X, tan) data. Only posterior probabilities passing the cut off used in cell type assignment (*L*_*N*_ ≥ 0.1, **Methods**) were plotted. **(c)** Spatial assignment of OBNBL1 cells. Spatial plots of posterior probabilities with assigned (color bar) or not assigned (lightgrey; only in 38X) barcode positions to the OBNBL1 cell type in all bin sizes (HDST, 5X and 38X). **(d)** Cell type assignment frequency. Percentage of barcode positions with a single cell type prediction out of total number of barcode positions. **(e)** Differentially expressed genes between all 63 cell types used in the prediction task on 5X “sc-like” bins. Shown is the scaled expression level (color bar) of most significant two genes (columns; bottom annotation) per cell type (columns; top annotation).

